# Genome Report: First whole genome sequence and assembly of the Ecuadorian brown-headed spider monkey (*Ateles fusciceps fusciceps*), a critically endangered species, using Oxford Nanopore Technologies

**DOI:** 10.1101/2023.08.29.555347

**Authors:** Gabriela Pozo, Martina Albuja Quintana, Lizbeth Larreátegui, Bernardo Gutiérrez, Nathalia Fuentes, Felipe Alfonso-Cortés, María de Lourdes Torres

## Abstract

The Ecuadorian brown-headed spider monkey (*Ateles fusciceps fusciceps*) is currently considered one of the most endangered primates in the world and is classified as critically endangered (IUCN). It faces multiple threats, the most significant one being habitat loss due to deforestation in western Ecuador. Genomic tools are key for the management of endangered species, but this requires a reference genome which until now was unavailable for *A. f. fusciceps*. The present study reports the first whole genome sequence and assembly of *A. f. fusciceps* generated using Oxford Nanopore long reads. DNA was extracted from a subadult male and libraries were prepared for sequencing following the Ligation Sequencing Kit SQK-LSK112 workflow. Sequencing was performed using a MinION Mk1C sequencer. The sequencing reads were processed to generate a genome assembly. Two different assemblers were used to obtain draft genomes using raw reads, of which the Flye assembly was found to be superior. The final assembly has a total length of 2.63 Gb and contains 3,861 contigs, with an N50 of 7,560,531 bp. The assembly was analyzed for annotation completeness based on primate ortholog prediction using a high-resolution database, and was found to be 84.3% complete, with a low number of duplicated genes indicating a precise assembly. The annotation of the assembly predicted 31,417 protein-coding genes, comparable to other mammal assemblies. A reference genome for this critically endangered species will allow researchers to gain insight into the genetics of its populations and thus aid conservation and management efforts of this vulnerable species.

## 1. Introduction

The brown-headed spider monkey (*Ateles fusciceps fusciceps*) is a neotropical primate inhabiting northwestern Ecuador (its presence in Colombia is uncertain). It is most commonly found below 1200 masl, but its altitudinal range can go as high as 2300 masl (Gallo-Viracocha et al., 2022). This subspecies plays an important role in the ecosystem as an effective seed disperser; its diet is composed mainly of ripe fruits (70-90%), which is key for the regeneration and maintenance of tree diversity in the forests it inhabits (Calle-Rendón et al., 2016; Gallo-Viracocha et al., 2022; Morelos Juarez et al., 2018).

*A. f. fusciceps* is a priority subject for conservation efforts worldwide, currently listed as one of the world’s 25 most endangered primates (Mittermeier et al., 2022) and cataloged as Critically Endangered by the IUCN (Moscoso et al., 2021). Anthropogenic factors are the main threats to *A. f. fusciceps* populations; as a large mammal with slow growth and reproduction rates, it is affected by the subsistence of hunting practices within indigenous communities, as well as poaching of infants for illegal wildlife trade. However, its most important threat is habitat loss. The Chocó region it inhabits in western Ecuador is a biodiversity hotspot (Myers et al., 2000) that requires immediate conservation action given that it has lost more than 80% of its original vegetation coverage (*Chocó-Darién-Western Ecuador: Chocó-Manabí Conservation Corridor Briefing Book*, 2005; Mittermeier et al., 1999; Myers et al., 2000; Sierra et al., 2021) .This has led to dramatic population decreases of several species in the region, including the brown-headed spider monkey (Moscoso et al., 2021). The current situation of *A. f. fusciceps* warrants a stronger focus on its conservation to prevent the extinction of the species.

Reductions of the numbers of individuals in brown-headed spider monkey populations make them susceptible to inbreeding depression and loss of genetic diversity through drift (Frankham, 2003; Rivera Román, 2017). These two processes reduce the species resilience to environmental change, thus increasing its vulnerability (Frankham, 2003; Romero, 2018). Whole Genome Sequencing (WGS) has been identified as a key tool to manage threatened species, as genomes from representative numbers of individuals can be used to make inferences on a population’s demographic history, inbreeding rates and past genetic bottlenecks, amongst other significant events (Taylor et al., 2022). For a critically endangered species like *A. f. fusciceps*, genomic population studies provide useful information regarding the species’ genetic diversity and population structure, which can assist with the design of adequate management regimes and conservation strategies such as those identified in the Conservation Action Plan for the Ecuadorian Primates (Tirira et al., 2018). Population genomic studies require a reference genome, which was not available for *A. f. fusciceps*.

Next Generation Sequencing has become more accessible in terms of costs and sequencing velocity. Nevertheless, limited resources in developing countries restrict the accessibility for usage and development of genomic tools (Helmy et al., 2016), especially for endangered species in the tropics (regions which harbor at least 50% of the planet’s biodiversity) (Brancalion et al., 2019). Oxford Nanopore sequencing has facilitated genomic research in developing countries with portable, low-cost sequencers that produce ultra-long reads and allow on-site sequencing (Lin et al., 2021). While only 1% of all threatened species have a published reference genome (Brandies et al., 2019), this could change as access to sequencing technologies increases. Given the overlap of high biodiversity and low accessibility to genomic tools, special emphasis and effort should be placed on genome sequencing projects of endangered species in developing nations.

In the present study we report the first whole genome sequence and assembly of *Ateles fusciceps fusciceps*, using long reads obtained through Oxford Nanopore Technologies.

## 2. Methods & Materials

### Sampling

The brown-headed spider monkey individual from which the sample was taken was a subadult male named Mishky, born in the Hacienda Jambelí Rescue Center (2°46’30.48”S 79°44’9.51”O) located in the Guayas province in southwestern Ecuador. In 2014, Proyecto Washu started an *ex situ* conservation program for the rehabilitation and welfare of this species. The Hacienda Jambelí population of *A. f. fusciceps* is currently considered the largest captive population in Ecuador with a total of 21 individuals: 8 adult males, 1 subadult male, 7 adult females, 1 subadult female, 1 juvenile female, and 3 juvenile males. This population is composed of individuals rescued from the illegal pet trade and others born in the rescue center, as is the case of Mishky.

Mishky was transported to the Tueri Wildlife Hospital (TUERI-USFQ) for medical examination due to injuries sustained while at the Hacienda Jambelí Rescue Center. A 5 ml blood sample was obtained by the TUERI-USFQ veterinarian staff and stored at -80°C in the Laboratorio de Biotecnología Vegetal – USFQ.

### Sequencing Methods and Preparation

#### a. DNA extraction

For DNA extraction, the DNeasy Blood and Tissue Kit (QIAGEN, Valencia, CA, USA) was used for 16 total reactions with minor modifications. For the final elution, 30 μl of ultrapure water was used to obtain a total elution of 60 μl after two elution steps. The final DNA quantification and quality was assessed with Qubit™ Fluorometric Quantitation and NanoDrop 2000.

#### b. Preparation of genomic libraries

The library construction protocol followed the workflow of the Ligation Sequencing Kit SQK-LSK112 (Oxford Nanopore Technologies), which comprises three sections. The process started with an average quantity of 2000 ng per reaction and resulted in a total of 14 libraries. After each section, the DNA concentration was quantified using Qubit™ Fluorometric Quantitation. The libraries were stored at 4°C awaiting sequencing.

#### c. Sequencing

Sequencing was carried out in a MinION Mk1C sequencer using six R10.4 flow cells. Each flow cell was used for three to four runs to generate a total of 21 sequencing runs (>24 h). The libraries that had a high DNA quantity (> 800 ng), were used for two sequencing runs. Similarly, depending on the final concentration of each library, 6, 7 or 12 μl of the sample was loaded to the flow cell, in order to sequence approximately 500 ng of DNA. The real-time base calling was executed with Guppy v5.1.13 (ONT) and the resulting output were raw fastq sequencing reads.

### Data processing

#### a. Initial processing of reads

The raw sequencing reads (.fastq) were first filtered according to quality scores using NanoFilt v2.3.0 (De Coster et al., 2018). Reads with quality scores below 7 were removed from the analysis (Feng et al., 2022; Halstead et al., 2021; Petersen et al., 2022). Adapters from filtered reads were then trimmed in Porechop v0.2.4 (Wick et al., 2017) and sequencing quality was analyzed in Nanoplot v. 1.20.0 (De Coster et al., 2018) for both individual sequencing runs and the complete dataset.

#### b. Assembly, mapping, polishing and scaffolding

Two different assemblers were used to obtain draft genomes using raw reads. First, SMARTdenovo v.1.0.0 (Liu et al., 2021) was used to assemble the obtained reads with the smartdenovo.pl script.

Raw reads were also assembled using Flye v 2.7.1 (Kolmogorov et al., 2019), selecting *nano-raw* as the type of input reads and with a specified genome size (g) of 2.6 Gb, based on the reported genome size of the closely related species *Ateles geoffroyi* (Shao, 2022).

Both *de novo* assembly drafts were mapped against the reference genome *of A. geoffroyi* using minimap2 v 2.24 (Li, 2018) to reorder the contigs generated in the assembly. The resulting mapped assemblies were then polished using Medaka v 1.7.2 (Oxford Nanopore Technologies, 2018). The *medaka_consensus* program was employed using the *r103_fast_g507* model.

#### c. Completeness and Quality Assessment of Genome Assembly

Genome assembly quality for both assemblies was evaluated with QUAST v 5.2.0 (Mikheenko et al., 2018) under default parameters. The reference genome of *A. geoffroyi* (Shao, 2022) was specified as the reference for comparison. BUSCO v 5.4.4 (Manni et al., 2021) was then run using the primates_odb10 database with 13,780 genes to evaluate genome completeness based on expected gene content; we provide statistics for complete, fragmented, duplicated, and missing BUSCOs.

#### d. Genome Annotation

The best assembly was selected based on the assembly statistics and BUSCO results, and that assembly was annotated. For genome annotation, a custom repeat library was first created *ab initio* for the assembled genome of *A. f. fusciceps* using RepeatModeler v 2.0.4 (Flynn et al., 2020). We applied the “LTRStruct” option for long terminal repeat (LTR) retroelement identification. Repetitive regions of the genome were identified and soft-masked by RepeatMasker v 4.0.7 (Smith et al., 2013) in Maker v 2.31.9 (Campbell et al., 2014). Contigs were then annotated with Maker v 2.31.9 (Campbell et al., 2014) in three consecutive rounds. In the first round, *ab initio* gene prediction algorithms were run with EST and protein evidence using the *est2genome* and *protein2genome* functions. Reference proteomes from four closely related primate species were gathered from the UniProt database (Bateman et al., 2021) to be used as protein evidence in Maker (*Sapajus apella: UP000504640. Callithrix jacchus*: UP000008225. *Saimiri boliviensis boliviensis*: UP000233220. *Aotus nancymaae*: UP000233020). EST data was obtained from the NCBI EST database from the most closely related species available (*Callithrix jacchus*). These initial predictions were then used to train the *ab initi*o gene predictor SNAP (Korf, 2004) and a second round of Maker was run using the hidden Markov model (HMM) from SNAP. Finally, a third round of annotation was run with SNAP. Protein and transcript fasta files and gff files generated along the three annotation rounds were then merged. To isolate the best supported gene models, InterProScan v 5.61 (Jones et al., 2014) was first run to identify conserved Pfam domains on the Maker predicted proteins. Using accessory scripts from Maker, gene models with AED values greater than 0.5 or lacking Pfam domains were then removed from the gff and fasta files. Finally, the agat_sp_statistics.pl script from the AGAT (Another Gff Analysis Toolkit) software was used to obtain annotation statistics (Dainat, 2020).

## 3. Results and Discussion

### *A. f. fusciceps* assembly

Oxford Nanopore Sequencing of *A. f. fusciceps* produced 55.95 Gb from 8.96 million reads. In order to calculate the coverage, we based our predicted genome size on the closely related species, *A. geofroyi* which is 2.6 Gb (Shao, 2022). This represents an estimated 21x coverage of the genome. In general, reads had a mean read length of 6.42 kb and a mean read quality score of 10.9 (Table 1). Since various reports of genome assemblies with Oxford Nanopore reads specify q7 as the threshold for acceptable read quality (Feng et al., 2022; Halstead et al., 2021; Petersen et al., 2022), only reads greater than or equal to q7 were used for posterior assembly.

**Table 1.** Sequencing statistics for the *Ateles fusciceps fusciceps* genome.

| Generated Bases | Read Count | Coverage | Mean Read Length | Mean Read Quality |
| --- | --- | --- | --- | --- |
| 55.95 Gb | 8,96 million | 21X | 6.42 Kb | 10.9 |

The assembly obtained with SMARTdenovo and later polished by Medaka had a total length of 2.58 Gb and contained 6,856 contigs (Table 2). It had an N50 size of 799,988 bp, an L50 of 985 and its largest contig was 5,164,154 bp. When mapped to the reference genome of the closely related *A. geoffroyi*, it had 567.9 mismatches per 100kbp. The Flye assembler alongside the Medaka polisher generated a primary assembly for *A. f. fusciceps* of 2.63 Gb containing 3,861 contigs with an N50 size of 7,560,531 bp (Table 2). The L50 for this assembly was 97 and the largest contig was 44,929,532 bp. In this case, when mapped to *A. geoffroyi* the assembly had 539.3 mismatches per 100 kbp.

**Table 2.** Statistics for the two obtained *A. f. fusciceps* assemblies using SMARTdenovo and Flye.

| Assembly | Total length | Contig number | N50 | Largest contig | L50 | # mismatches per 100 kbp |
| --- | --- | --- | --- | --- | --- | --- |
| SMARTdenovo | 2,586,824,631 | 6,856 | 799,988 | 5,164,154 | 985 | 567.9 |
| Flye | 2,635,867,907 | 3,861 | 7,560,531 | 44,929,532 | 97 | 539.3 |

The Flye assembly is superior to the SMARTdenovo assembly in all analyzed statistics (Table 2). It has a total length similar to the genome size of the closely related *A. geoffroyi* (2.68 Gb) (Shao, 2022) and less mismatches per 100 kbp when compared to this genome. It is much less fragmented, with 3,861 contigs compared to 6,856 in the SMARTdenovo assembly. Furthermore, according to the L50, 50% of the *A. f. fusciceps* genome is represented in 97 contigs in the Flye assembly, and in 985 contigs in the the SMARTdenovo assembly, proving once again that the SMARTdenovo assembly is less continuous. The Flye assembly also has a much higher N50 and largest contig size; 50% of the contigs possess a size equal to or longer than 7.56 Mb (Alhakami et al., 2017) which is remarkable since primate species have very large genomes and first assemblies normally produce contig N50 lengths shorter than 100 kbp (Jayakumar et al., 2021). Furthermore, the largest contig size of the Flye assembly is 44.9 Mb, almost the size of a human chromosome (Brown, 2002).

Both genome assemblies were analyzed for annotation completeness based on primate ortholog prediction. The gene database used, primates_odb10, comprises 25 primate genomes and 13,780 genes, and is categorized as a high-resolution database which provides a high level of confidence for genome completeness evaluations (Simão et al., 2015; Waterhouse et al., 2018). For the SMARTdenovo assembly we obtained 10,602 (76.9%) complete BUSCOs, of which 10,384 are single copy (75.4%) and 218 (1.6%) are duplicated (Fig. 1). There were 2,436 (17,7%) missing BUSCOs and 742 (5.4%) fragmented BUSCOs.

**Figure 1.**
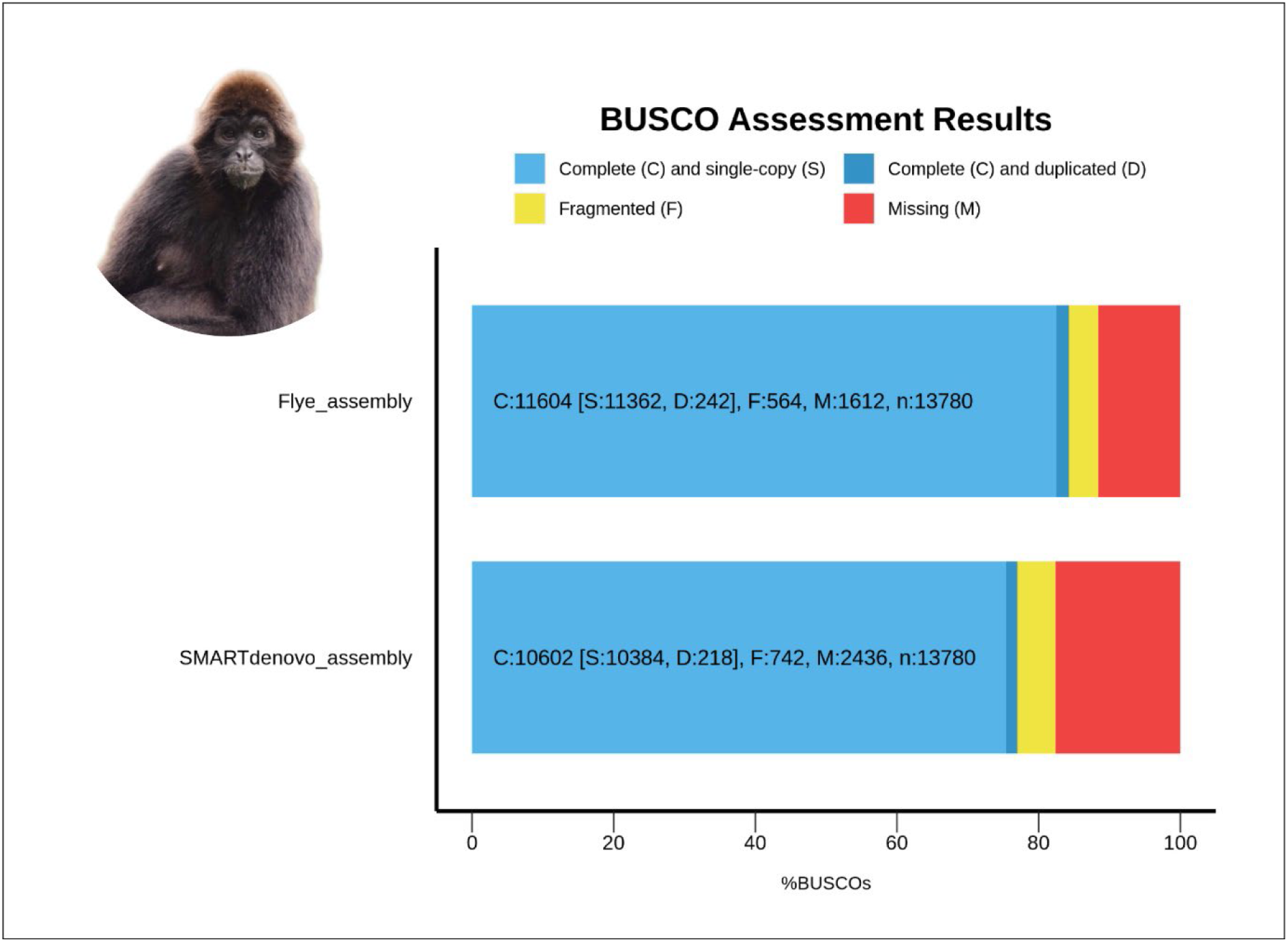
BUSCO results for SMARTdenovo and Flye *A.f.fusciceps* assemblies. Complete and Single-Copy, Complete and Duplicated, Fragmented and Missing BUSCOS are presented. Primates_odb10, containing 13,780 genes, was used as the reference database.

When analyzing the Flye assembly, the BUSCO results improved: we obtained more single-copy complete BUSCOs and less missing or fragmented BUSCOs. Specifically, we obtained 11,604 (84.3%) complete BUSCOs, of which 11,362 (82.5%) are single copy and 242 (1.8%) are duplicated (Fig. 1). The high number of complete BUSCOs (84.3%) and the low number of duplicated genes indicate a good level of genome completeness and a precise assembly (Manni et al., 2021; Simão et al., 2015). Regarding the remaining 15.7% of BUSCOs, 564 (4.1%) are fragmented and 1,612 (11.6%) are missing. Technical limitations in gene prediction can inflate the proportions of missing and fragmented BUSCOs, when working with large genomes such as that of *A. f. fusciceps* (Manni et al., 2021). Additionally, ONT sequences have error rates of 10% to 30% that are mainly composed of indels (Morisse et al., 2021). However, while the assembly could be improved, the results indicate an overall good quality of the Flye assembly.

Due to the fact that the Flye assembly has better assembly statistics and a more complete annotation, this is the one we selected for further analyses and the one that is reported in this publication. Our *A. f. fusciceps* assembly was compared to that of the closely related *A. geoffroyi* (GCA_023783555.1) (Table 3). This contig-level assembly of *A. geoffroyi* has a total length of 2.68 Gb in 2,732 contigs with a N50 size of 29,212,752 bp and a GC content of 40.75%. The values for coverage, contig number an N50 size for both assemblies were significantly different. However, considering that the range of genome size variation among primates is small (Fantini et al., 2016) and that primate genomes GC-contents are remarkably consistent (Qi et al., 2016), the similar values for total length and GC (%) clearly show that this primary genome assembly of *A. f. fusciceps* is adequate, while the differences in coverage, contig number and N50 suggest there is room for improvement.

**Table 3.** Assembly statistics for the *A. f. fusciceps* genome compared to the closely related *A. geoffroyi* assembly.

| Genome | Total length | Coverage | Contig number | N50 | GC (%) |
| --- | --- | --- | --- | --- | --- |
| *A. fusciceps* | 2,635,863,941 | 21x | 3,861 | 7,560,531 | 40.86 |
| *A. geoffroyi* | 2,683,028,796 | 56.87x | 2,723 | 29,212,752 | 40.75 |

### Genome Annotation

The annotation of the *A. f. fusciceps* assembly in Maker predicted 35,809 protein-coding genes, 88% (31,417) with an AED value < 0.5 (Table 4), indicating good protein and transcript evidence support and reasonable quality of the annotation (Saenko et al., 2021; Sork et al., 2016). AED values closer to 0 generally show greater agreement between the annotation and protein/transcript evidence while AED values closer to 1 reveal little to no support for the resulting annotation (Eilbeck et al., 2009); which is why all gene models with AED values > 0.5 were filtered out of the final annotation.

**Table 4.**
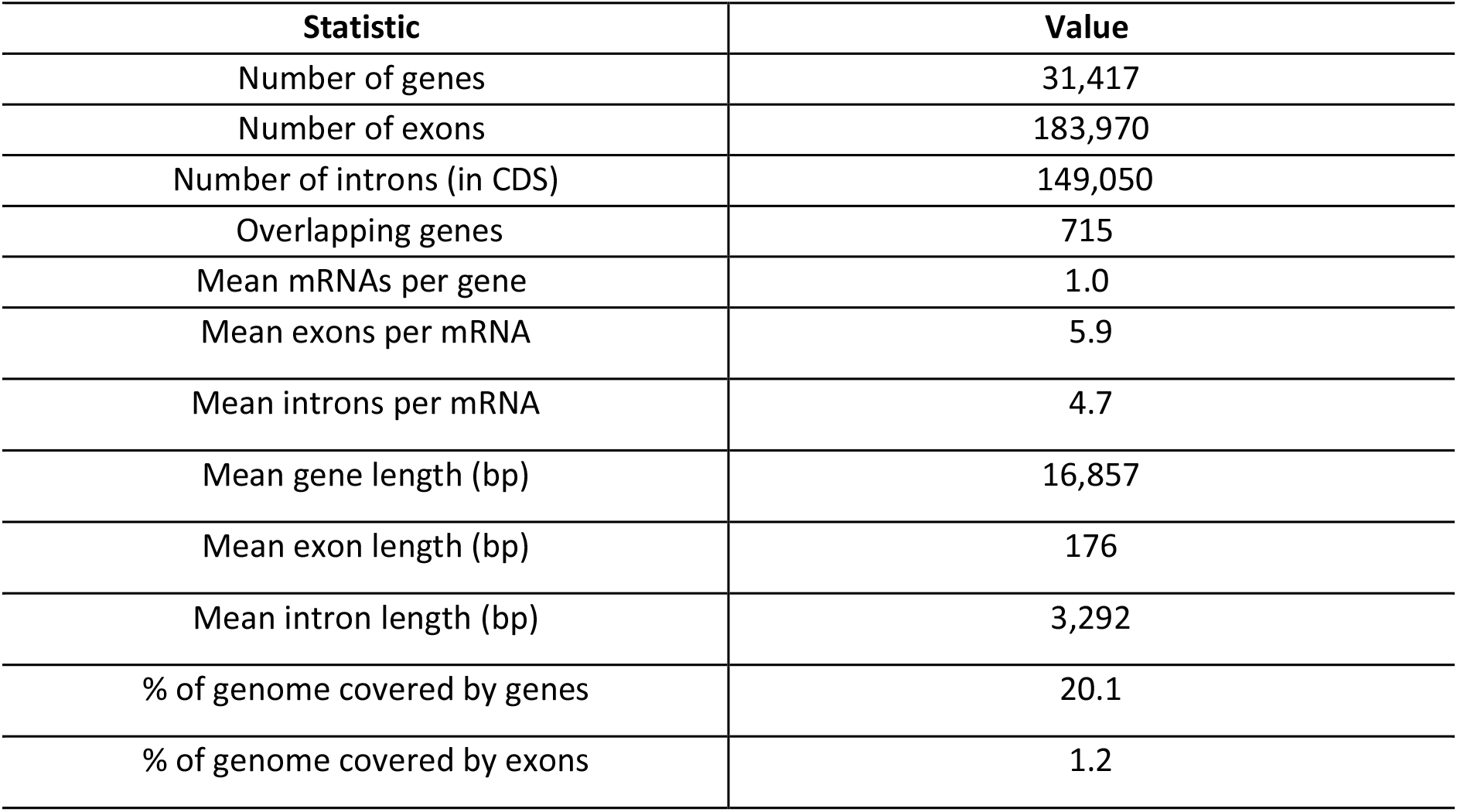
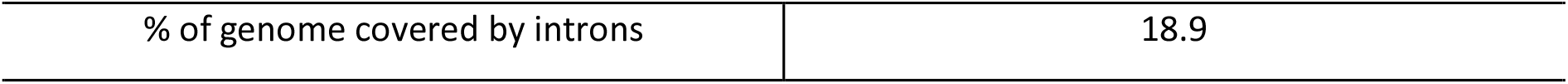
Summary Statistics of the annotated genome (AED<0.5) of *A. f. fusciceps*.

| Statistic | Value |
| --- | --- |
| Number of genes | 31,417 |
| Number of exons | 183,970 |
| Number of introns (in CDS) | 149,050 |
| Overlapping genes | 715 |
| Mean mRNAs per gene | 1.0 |
| Mean exons per mRNA | 5.9 |
| Mean introns per mRNA | 4.7 |
| Mean gene length (bp) | 16,857 |
| Mean exon length (bp) | 176 |
| Mean intron length (bp) | 3,292 |
| % of genome covered by genes | 20.1 |
| % of genome covered by exons | 1.2 |
| % of genome covered by introns | 18.9 |

The resulting 31,417 protein-coding genes of *A. f. fusciceps* are comparable to what other mammal genome assemblies have reported like the case of the lowland anoa (*Bubalus depressicornis*) with 32,393 predicted protein-coding genes (Porrelli et al., 2022). Nonetheless, gene count is slightly higher than expected when compared to the 22,027 protein-coding genes predicted for *Callithrix jacchus* (GCA_011100555.1) (Warren et al., 2009) and the 20,350 protein-coding genes for *Sapajus apella* (GCF_009761245.1) (Culibrk et al., 2019); both closely related primate species of *A. f. fusciceps*. In general, eukaryotic genomes have around 15,000–25,000 protein coding genes (Cantarel et al., 2008) with the human genome (a primate species) reporting around ∼19,100 genes (Piovesan et al., 2019). The overestimation of the protein-coding genes could be explained by ONT’s long read accuracy limitations compared to other sequencing technologies (Rang et al., 2018), though the resulting annotation of our genome still shows an accurate prediction. Additionally, since only soft masking was used for repeat masking during MAKER annotation, it is possible that repetitive regions were misconceived as putative genes (Saenko et al., 2021), increasing the predicted number of coding sequences.

Furthermore, the annotation of the *A. f. fusciceps* genome predicted a mean gene length of 16,857 bp (Table 4); a length comparably smaller to what has been reported for other closely related primate species, with mean gene lengths of around ∼40,000 bp (Culibrk et al., 2019; *Saimiri Boliviensis Boliviensis Breed Bolivian Squirrel Monkey Isolate - Nucleotide - NCBI*, n.d.; Warren et al., 2009). The same pattern is evident when we compare mean intron length (3,292 bp) and mean exon length (176 bp). These differences can likely be attributed to the level of fragmentation of our genome and the inaccurate prediction of genomic features in repetitive regions. This is expected since around 50% of a primate genome is covered by repetitive elements (Rogers & Gibbs, 2014), making the annotation of other genomic features a challenging task (Okazaki & Hume, 2003). Nonetheless, differences in genomic feature predictions between closely related species have been reported in other reference genomes (Jiang et al., 2022; Kaur et al., 2023) and could be attributed to the sequencing technology used and the level of genome fragmentation.

### Importance of reference genome

Numerous studies have established the importance of genomic data to understand the evolutionary history of a species and to develop appropriate conservation and management strategies (Kenny et al., 2020; Kleinman-Ruiz et al., 2017; Nong et al., 2021; Pfenninger et al., 2021; Saremi et al., 2019). Whole-genome sequencing (WGS) leads to a better understanding of the biology of a species and provides insights into fundamental processes that shape their evolution (Ryder, 2005), and its application can provide important and accurate information about its demographic history, admixture, introgression, recombination, linkage disequilibrium, genomic regions evolving under selective pressures and other evolutionary processes (Theissinger et al., 2023). For critically endangered species like the brown-headed spider monkey, genomic approaches are even more valuable due to the scarcity of samples for genetic studies; therefore, WGS maximizes the information that researchers can harness from each sample. However, in order to be able to generate and fully take advantage of this information, a reference genome is required (Theissinger et al., 2023).

Species under such conservation threats face a dire need for conservation actions to reverse their declining population trends. Currently, Proyecto Washu is deepening the understanding of the brown-headed spider monkey’s behavior and ecology through observational studies of a population of spider monkeys living in a highly fragmented landscape. The sequencing of its genome provides an opportunity to improve its conservation through the development of population-level studies to evaluate their genetic diversity and gene flow. Moreover, genetic population studies may allow us to better differentiate its populations, perform identification of individuals and kinship patterns, evaluate the dispersion and migration of individuals, and identify and prioritize biological corridors through which monkey populations move. Biological corridors prevent the isolation of populations in closed forest fragments which reduces inbreeding and helps to maintain genetic diversity in the area (Haddad et al., 2015; Kirchner et al., 2003).

While major progress has been made on animal genome sequencing in the last 25 years, significant gaps and biases remain in geographic and taxonomic representation resulting in an improper depiction of the global genetic pool (Hotaling et al., 2021). Ecuador, for instance, has a limited record of genetic and genomic research (Zambrano-Mila et al., 2019) despite its sizable biodiversity (Celi & Villamarín, 2020). This is a multifaceted issue resulting from the lack of sequencing platforms, training in genome data analysis and research costs (Hotaling et al., 2021). This makes outsourcing a popular alternative to generate genomic sequences, despite the limitations of using third-party service providers (Helmy et al., 2016). A feasible pathway to democratize sequencing efforts and to involve developing countries is through the usage of portable sequencing devices such as the Oxford Nanopore Technologies MinION, as applied in this study. This is a time- and cost-efficient technology for the assembly of all genome sizes (Wang et al., 2021) which operates on standard computing resources. Its long read length and portability enables the use of these devices in basic research (e.g., assembly of preliminary non-model organism genomes), clinical usage and on-site applications (Wang et al., 2021). Due to its ease of use and convenience, the current report represents an initial sequencing project which will be further extended to other underrepresented Ecuadorian mammals. We expect that this and similar efforts will generate critical information for future genomic studies directed towards conservation and management efforts.

## 4. Conclusions

The brown-headed spider monkey (*Ateles fusciceps fusciceps*) is a critically endangered primate species, facing multiple threats such as habitat loss and hunting, emphasizing the urgent need for conservation efforts. Whole Genome Sequencing (WGS) has been identified as a crucial tool for managing threatened species. Here, we present the first whole genome sequence and assembly of *A. f. fusciceps* using long reads obtained through Oxford Nanopore Technologies, which resulted in a good quality assembly. The genomic insights gained from this study provide valuable information which can lead to the development of tools for the conservation of *A. f. fusciceps*. Moreover, the pipelines used in this study can serve as a foundation for sequencing and assembling genomes of other endangered species in developing nations, ultimately aiding in the preservation of global biodiversity.

## 5. Data Availability Statement

The genome assembly and annotation are available at NCBI with accession number (SUB13795553).

## 6. Acknowledgements

We thank Carolina Sáenz and TUERI-USFQ for their valuable aid in obtaining the sample of the sequenced individual. We also thank the members of the Plant Biotechnology Laboratory (USFQ) for their input and help during this research.

Genetic data for the specimen was obtained under the Genetic Resources Permit Number: MAE-DNB-CM-2019-0126 granted to INABIO by Ministerio del Ambiente, Agua y Transición Ecológica in Ecuador, in accordance with Ecuadorian law.

## References

Alhakami, H., Mirebrahim, H., & Lonardi, S. (2017). A comparative evaluation of genome assembly reconciliation tools. Genome Biology, 18(1). 10.1186/s13059-017-1213-3

Bateman, A., Martin, M. J., Orchard, S., Magrane, M., Agivetova, R., Ahmad, S., Alpi, E., Bowler-Barnett, E. H., Britto, R., Bursteinas, B., Bye-A-Jee, H., Coetzee, R., Cukura, A., da Silva, A., Denny, P., Dogan, T., Ebenezer, T. G., Fan, J., Castro, L. G., … Teodoro, D. (2021). UniProt: the universal protein knowledgebase in 2021. Nucleic Acids Research, 49(D1), D480–D489. 10.1093/NAR/GKAA1100

Brancalion, P. H. S., Niamir, A., Broadbent, E., Crouzeilles, R., Barros, F. S. M., Zambrano, A. M. A., Baccini, A., Aronson, J., Goetz, S., Reid, J. L., Strassburg, B. B. N., Wilson, S., & Chazdon, R. L. (2019). APPLIEDECOLOGY Global restoration opportunities in tropical rainforest landscapes. In Sci. Adv (Vol. 5). https://www.science.org

Brandies, P., Peel, E., Hogg, C. J., & Belov, K. (2019). The Value of Reference Genomes in the Conservation of Threatened Species. Genes 2019, Vol. 10, Page 846, 10(11), 846. 10.3390/GENES10110846

Brown, T. A. (2002). Chapter 1, The Human Genome. In Genomes (2nd Editio). Wiley-Liss.

Calle-Rendón, B. R., Peck, M., Bennett, S. E., Morelos-Juarez, C., & Alfonso, F. (2016). Comparison of forest regeneration in two sites with different primate abundances in Northwestern Ecuador. Revista de Biología Tropical, 64(2), 493–506.

Campbell, M. S., Holt, C., Moore, B., & Yandell, M. (2014). Genome Annotation and Curation Using MAKER and MAKER-P. Current Protocols in Bioinformatics, 48, 4.11.1-4.11.39. 10.1002/0471250953.BI0411S48

Cantarel, B. L., Korf, I., Robb, S. M. C., Parra, G., Ross, E., Moore, B., Holt, C., Sánchez Alvarado, A., & Yandell, M. (2008). MAKER: An easy-to-use annotation pipeline designed for emerging model organism genomes. Genome Research, 18(1), 188–196. 10.1101/gr.6743907

Celi, J. E., & Villamarín, F. (2020). Freshwater ecosystems of Mainland Ecuador: diversity, issues and perspectives. Acta Limnologica Brasiliensia, 32. 10.1590/s2179-975x3220

Chocó-Darién-Western Ecuador: Chocó-Manabí Conservation Corridor Briefing Book. (2005). https://www.cepf.net/sites/default/files/final.chocodarienwesternecuador.chocomanabi.briefingbook.

Culibrk, L., Leelakumari, S., Tse, K., Cheng, D., Chuah, E., Kirk, H., Pandoh, P., Troussard, A., Zhao, Y., Mungall, A., Moore, R., Marra, M. A. ., Sinclair-Smith, T., & Jones, S. J. M. (2019). The genome of the Tufted Capuchin (Sapajus apella). https://www.ncbi.nlm.nih.gov/nuccore/WRPQ00000000.1/

Dainat, J. (2020). Another Gff Analysis Toolkit to handle annotations in any GTF/GFF format (v0.8.0). Zenodo. 10.5281/zenodo.3552717

De Coster, W., D’Hert, S., Schultz, D. T., Cruts, M., & Van Broeckhoven, C. (2018). NanoPack: visualizing and processing long-read sequencing data. Bioinformatics, 34(15), 2666–2669. 10.1093/BIOINFORMATICS/BTY149

Eilbeck, K., Moore, B., Holt, C., & Yandell, M. (2009). Quantitative measures for the management and comparison of annotated genomes. BMC Bioinformatics, 10(1), 67. 10.1186/1471-2105-10-67

Fantini, L. I., Jeffery, N. W., Pierossi, P., Ryan Gregory, T., & Nieves, M. (2016). Qualitative and quantitative analysis of the genomes and chromosomes of spider monkeys (Primates: Atelidae). http://www.genomesize.com

Feng, L., Lin, H., Kang, M., Ren, Y., Yu, X., Xu, Z., Wang, S., Li, T., Yang, W., & Hu, Q. (2022). A chromosome-level genome assembly of an alpine plant Crucihimalaya lasiocarpa provides insights into high-altitude adaptation. DNA Research, 29(1). 10.1093/DNARES/DSAC004

Flynn, J. M., Hubley, R., Goubert, C., Rosen, J., Clark, A. G., Feschotte, C., & Smit, A. F. (2020). RepeatModeler2 for automated genomic discovery of transposable element families. Proceedings of the National Academy of Sciences of the United States of America, 117(17), 9451–9457. 10.1073/PNAS.1921046117/SUPPL_FILE/PNAS.1921046117.SAPP.PDF

Frankham, R. (2003). Genetics and conservation biology. Comptes Rendus Biologies, 326(SUPPL. 1), 22–29. 10.1016/S1631-0691(03)00023-4

Gallo-Viracocha, F., Urgilés-Verdugo, C., Fuentes, N., Alfonso-Cortes, F., Zurita-Arthos, L., Torres, T. C., & Tirira, D. G. (2022). Distribution, conservation, and vulnerability to climate change of the Ecuadorian Brown-headed Spider Monkey (Primates: Atelidae). Mammalia Aequatorialis, 4. https://mamiferosdelecuador.com/mammalia-aequatorialis/index.php/boletin/article/view/50

Haddad, N. M., Brudvig, L. A., Clobert, J., Davies, K. F., Gonzalez, A., Holt, R. D., Lovejoy, T. E., Sexton, J. O., Austin, M. P., Collins, C. D., Cook, W. M., Damschen, E. I., Ewers, R. M., Foster, B. L., Jenkins, C. N., King, A. J., Laurance, W. F., Levey, D. J., Margules, C. R., … Townshend, J. R. (2015). Habitat fragmentation and its lasting impact on Earth’s ecosystems. Science Advances, 1(2). 10.1126/sciadv.1500052

Halstead, M. M., Islas-Trejo, A., Goszczynski, D. E., Medrano, J. F., Zhou, H., & Ross, P. J. (2021). Large-Scale Multiplexing Permits Full-Length Transcriptome Annotation of 32 Bovine Tissues From a Single Nanopore Flow Cell. Frontiers in Genetics, 12, 621. 10.3389/FGENE.2021.664260/BIBTEX

Helmy, M., Awad, M., & Mosa, K. A. (2016). Limited resources of genome sequencing in developing countries: Challenges and solutions. Applied & Translational Genomics, 9, 15–19. 10.1016/J.ATG.2016.03.003

Hotaling, S., Kelley, J. L., & Frandsen, P. B. (2021). Toward a genome sequence for every animal: Where are we now? Proceedings of the National Academy of Sciences of the United States of America, 118(52). 10.1073/PNAS.2109019118/-/DCSUPPLEMENTAL

Jayakumar, V., Nishimura, O., Kadota, M., Hirose, N., Sano, H., Murakawa, Y., Yamamoto, Y., Nakaya, M., Tsukiyama, T., Seita, Y., Nakamura, S., Kawai, J., Sasaki, E., Ema, M., Kuraku, S., Kawaji, H., & Sakakibara, Y. (2021). Chromosomal-scale de novo genome assemblies of Cynomolgus Macaque and Common Marmoset. Scientific Data, 8(1). 10.1038/s41597-021-00935-6

Jiang, F., Wang, S., Wang, H., Wang, A., Xu, D., Liu, H., Yang, B., Yuan, L., Lei, L., Chen, R., Li, W., & Fan, W. (2022). A chromosome-level reference genome of a Convolvulaceae species Ipomoea cairica. G3 Genes|Genomes|Genetics, 12(9). 10.1093/G3JOURNAL/JKAC187

Jones, P., Binns, D., Chang, H.-Y., Fraser, M., Li, W., McAnulla, C., McWilliam, H., Maslen, J., Mitchell, A., Nuka, G., Pesseat, S., Quinn, A. F., Sangrador-Vegas, A., Scheremetjew, M., Yong, S.-Y., Lopez, R., & Hunter, S. (2014). InterProScan 5: genome-scale protein function classification. Bioinformatics, 30(9), 1236–1240. 10.1093/bioinformatics/btu031

Kaur, S., Stinson, S. A., & diCenzo, G. C. (2023). Whole genome assemblies of Zophobas morio and Tenebrio molitor. G3 Genes|Genomes|Genetics, 13(6), 79. 10.1093/G3JOURNAL/JKAD079

Kenny, N. J., Francis, W. R., Rivera-Vicéns, R. E., Juravel, K., de Mendoza, A., Díez-Vives, C., Lister, R., Bezares-Calderón, L. A., Grombacher, L., Roller, M., Barlow, L. D., Camilli, S., Ryan, J. F., Wörheide, G., Hill, A. L., Riesgo, A., & Leys, S. P. (2020). Tracing animal genomic evolution with the chromosomal-level assembly of the freshwater sponge Ephydatia muelleri. Nature Communications, 11(1). 10.1038/s41467-020-17397-w

Kirchner, F., Ferdy, J.-B., Andalo, C., Colas, B., & Moret, J. (2003). Role of Corridors in Plant Dispersal: an Example with the Endangered Ranunculus nodif lorus. Conservation Biology, 17(2), 401–410. 10.1046/j.1523-1739.2003.01392.x

Kleinman-Ruiz, D., Martínez-Cruz, B., Soriano, L., Lucena-Perez, M., Cruz, F., Villanueva, B., Fernández, J., & Godoy, J. A. (2017). Novel efficient genome-wide SNP panels for the conservation of the highly endangered Iberian lynx. BMC Genomics, 18(1). 10.1186/s12864-017-3946-5

Kolmogorov, M., Yuan, J., Lin, Y., & Pevzner, P. A. (2019). Assembly of long, error-prone reads using repeat graphs. Nature Biotechnology 2019 37:5, 37(5), 540–546. 10.1038/s41587-019-0072-8

Korf, I. (2004). Gene finding in novel genomes. BMC Bioinformatics, 5(59). 10.1186/1471-2105-5-59

Li, H. (2018). Minimap2: pairwise alignment for nucleotide sequences. Bioinformatics, 34(18), 3094–3100. 10.1093/bioinformatics/bty191

Lin, B., Hui, J., & Mao, H. (2021). Nanopore technology and its applications in gene sequencing. In Biosensors (Vol. 11, Issue 7). MDPI. 10.3390/bios11070214

Liu, H., Wu, S., Li, A., & Ruan, J. (2021). SMARTdenovo: a de novo assembler using long noisy reads. Gigabyte, 2021, 1–9. 10.46471/gigabyte.15

Manni, M., Berkeley, M. R., Seppey, M., & Zdobnov, E. M. (2021). BUSCO: Assessing Genomic Data Quality and Beyond. Current Protocols, 1(12), e323. 10.1002/CPZ1.323

Mikheenko, A., Prjibelski, A., Saveliev, V., Antipov, D., & Gurevich, A. (2018). Versatile genome assembly evaluation with QUAST-LG. Bioinformatics, 34(13), i142–i150. 10.1093/BIOINFORMATICS/BTY266

Mittermeier, R. A., Myers, N., Mittermeier, C. G., & Robles Gil, P. (1999). Hotspots: Earth’s biologically richest and most endangered terrestrial ecoregions. Hotspots: Earth’s Biologically Richest and Most Endangered Terrestrial Ecoregions.

Mittermeier, R. A., Reuter, K. E., Rylands, A. B., Jerusalinsky, L., Schwitzer, C., Strier, K. B., Ratsimbazafy, J., & Humle, T. (2022). Primates in Peril: The World’s 25 Most Endangered Primates 2022–2023.

Morelos Juarez, C., Tapia, A., Cervera, L., Alfonso-Cortez, F., Fuentes, N., Araguillin, E., Zapato-Ríos, G., Spaan, D., & Peck, M. R. (2018). Distribución actual, ecología y estrategias para la conservación de un primate críticamente amenazado (Ateles fusciceps fusiceps) en el Ecuador. La Primatología En Latino-América 2 – A Primatologia Na America Latina 2. Tomo II Costa Rica-Venezuela, 471–480. http://www.internationalprimatologicalsociety.org/docs/PrimLatam2-T.II-C.R-Vzla.pdf

Morisse, P., Marchet, C., Limasset, A., Lecroq, T., & Lefebvre, A. (2021). Scalable long read selfcorrection and assembly polishing with multiple sequence alignment. Scientific Reports |, 11, 761. 10.1038/s41598-020-80757-5

Moscoso, P., Link, A., de la Torra, S., Shanee, S., & Cortes-Ortíz, L. (2021). Ateles fusciceps ssp. fusciceps (Brown-headed Spider Monkey). The IUCN Red List of Threatened Species 2021: E.T39922A191687911. https://www.iucnredlist.org/species/39922/191687911

Myers, N., Mittermeler, R. A., Mittermeler, C. G., Da Fonseca, G. A. B., & Kent, J. (2000). Biodiversity hotspots for conservation priorities. Nature 2000 403:6772, 403(6772), 853–858. 10.1038/35002501

Nong, W., Qu, Z., Li, Y., Barton-Owen, T., Wong, A. Y. P., Yip, H. Y., Lee, H. T., Narayana, S., Baril, T., Swale, T., Cao, J., Chan, T. F., Kwan, H. S., Ngai, S. M., Panagiotou, G., Qian, P. Y., Qiu, J. W., Yip, K. Y., Ismail, N., … Hui, J. H. L. (2021). Horseshoe crab genomes reveal the evolution of genes and microRNAs after three rounds of whole genome duplication. Communications Biology, 4(1). 10.1038/s42003-020-01637-2

Okazaki, Y., & Hume, D. A. (2003). A Guide to the Mammalian Genome: Figure 1. Genome Research, 13(6b), 1267–1272. 10.1101/gr.1445603

Petersen, C., Sørensen, T., Westphal, K. R., Fechete, L. I., Sondergaard, T. E., Sørensen, J. L., & Nielsen, K. L. (2022). High molecular weight DNA extraction methods lead to high quality filamentous ascomycete fungal genome assemblies using Oxford Nanopore sequencing. Microbial Genomics, 8(4), 000816. 10.1099/MGEN.0.000816/CITE/REFWORKS

Pfenninger, M., Reuss, F., Kiebler, A., Schönnenbeck, P., Caliendo, C., Gerber, S., Cocchiararo, B., Reuter, S., Blüthgen, N., Mody, K., Mishra, B., Bálint, M., Thines, M., & Feldmeyer, B. (2021). Genomic basis for drought resistance in european beech forests threatened by climate change. ELife, 10. 10.7554/eLife.65532

Piovesan, A., Antonaros, F., Vitale, L., Strippoli, P., Pelleri, M. C., & Caracausi, M. (2019). Human protein-coding genes and gene feature statistics in 2019. BMC Research Notes, 12(1). 10.1186/S13104-019-4343-8

Porrelli, S., Gerbault-Seureau, M., Rozzi, R., Chikhi, R., Curaudeau, M., Ropiquet, A., & Hassanin, A. (2022). Draft genome of the lowland anoa (Bubalus depressicornis) and comparison with buffalo genome assemblies (Bovidae, Bubalina). G3 Genes|Genomes|Genetics, 12(11). 10.1093/g3journal/jkac234

Qi, W.-H., Yan, C., Li, W.-J., Jiang, X.-M., Li, G.-Z., Zhang, X.-Y., Hu, T.-Z., Li, J., & Yue, B.-S. (2016). Distinct patterns of simple sequence repeats and GC distribution in intragenic and intergenic regions of primate genomes. Aging, 8(11), 2635–2654. 10.18632/aging.101025

Rang, F. J., Kloosterman, W. P., & de Ridder, J. (2018). From squiggle to basepair: Computational approaches for improving nanopore sequencing read accuracy. Genome Biology, 19(1), 1–11. 10.1186/S13059-018-1462-9/FIGURES/3

Rivera Román, E. S. (2017). Filogeografía del mono araña de cabeza café (Ateles fusciceps fusciceps) en el Ecuador [Universidad Central del Ecuador]. http://www.dspace.uce.edu.ec/bitstream/25000/11043/1/T-UCE-0016-009.pdf

Rogers, J., & Gibbs, R. A. (2014). Comparative primate genomics: emerging patterns of genome content and dynamics. Nature Reviews. Genetics, 15(5), 347. 10.1038/NRG3707

Romero, V. (2018). Ateles fusciceps. In J. Brito, M. A. Camacho, V. Romero, & A. F. Vallejo (Eds.), Mamíferos del Ecuador (Version 20). Museo de Zoología, Pontificia Universidad Católica del Ecuador. https://bioweb.bio/faunaweb/mammaliaweb/FichaEspecie/Atelesfusciceps

Ryder, O. A. (2005). Conservation genomics: applying whole genome studies to species conservation efforts. Cytogenetic and Genome Research, 108(1–3), 6–15. 10.1159/000080796

Saenko, S. V, Groenenberg, D. S. J., Davison, A., & Schilthuizen, M. (2021). The draft genome sequence of the grove snail Cepaea nemoralis. G3 Genes|Genomes|Genetics, 11(2). 10.1093/g3journal/jkaa071

Saimiri boliviensis boliviensis breed Bolivian squirrel monkey isolate -Nucleotide -NCBI. (n.d.). Retrieved April 28, 2023, from https://www.ncbi.nlm.nih.gov/nuccore/1955098031

Saremi, N. F., Supple, M. A., Byrne, A., Cahill, J. A., Coutinho, L. L., Dalén, L., Figueiró, H. V., Johnson, W. E., Milne, H. J., O’Brien, S. J., O’Connell, B., Onorato, D. P., Riley, S. P. D., Sikich, J. A., Stahler, D. R., Villela, P. M. S., Vollmers, C., Wayne, R. K., Eizirik, E., … Shapiro, B. (2019). Puma genomes from North and South America provide insights into the genomic consequences of inbreeding. Nature Communications, 10(1). 10.1038/s41467-019-12741-1

Shao, Y. (2022). Ateles geoffroyi isolate KIZ-2021_1, whole genome shotgun sequencing p -Nucleotide -NCBI. Phylogenomic Analyses Provide Insights into Primate Genomic and Phenotypic Evolution. https://www.ncbi.nlm.nih.gov/nuccore/JAKFHY000000000.1

Sierra, R., Calva, O., & Guevara, A. (2021). La Deforestación en el Ecuador, 1990-2018. Factores promotores y tendencias recientes. https://www.proamazonia.org/wp-content/uploads/2021/06/Deforestación_Ecuador_com2.pdf

Simão, F. A., Waterhouse, R. M., Ioannidis, P., Kriventseva, E. V., & Zdobnov, E. M. (2015). BUSCO: assessing genome assembly and annotation completeness with single-copy orthologs. Bioinformatics, 31(19), 3210–3212. 10.1093/bioinformatics/btv351

Smith, A., Hubley, R., & Green, P. (2013). RepeatMasker Open-4.0. RepeatMasker Open-4.0.

Sork, V. L., Fitz-Gibbon, S. T., Puiu, D., Crepeau, M., Gugger, P. F., Sherman, R., Stevens, K., Langley, C. H., Pellegrini, M., & Salzberg, S. L. (2016). First Draft Assembly and Annotation of the Genome of a California Endemic Oak Quercus lobata Née (Fagaceae). G3 Genes|Genomes|Genetics, 6(11), 3485–3495. 10.1534/g3.116.030411

Taylor, R. S., Manseau, M., Redquest, B., Sonesinh Keobouasone, ·, Patrick Gagné, ·, Martineau, C., & Wilson, P. J. (2022). Whole genome sequences from non-invasively collected caribou faecal samples. 14, 53–68. 10.1007/s12686-021-01235-2

Theissinger, K., Fernandes, C., Formenti, G., Bista, I., Berg, P. R., Bleidorn, C., Bombarely, A., Crottini, A., Gallo, G. R., Godoy, J. A., Jentoft, S., Malukiewicz, J., Mouton, A., Oomen, R. A., Paez, S., Zhang, G., Jarvis, E. D., Bálint, M., Ciofi, C., … European, T. (2023). 20 Christophe Pampoulie, 21 María J. Ruiz-López, 16,22 Simona Secomandi, 15 Hannes Svardal. 19. 10.1016/j.tig.2023.01.005

Tirira, D. G., de la Torre, S., & Ríos, G. Z. (2018). Plan de acción para Estudio, de los primates del Ecuador.

Wang, Y., Zhao, Y., Bollas, A., Wang, Y., & Au, K. F. (2021). Nanopore sequencing technology, bioinformatics and applications. Nature Biotechnology, 39(11), 1348–1365. 10.1038/s41587-021-01108-x

Warren, W., Ye, L., Minx, P., Worley K., Gibbs, R., & Wilson, R. K. (2009). Proteomes_Callithrix jacchus (White-tufted-ear marmoset).

Waterhouse, R. M., Seppey, M., Simão, F. A., Manni, M., Ioannidis, P., Klioutchnikov, G., Kriventseva, E. V, & Zdobnov, E. M. (2018). BUSCO Applications from Quality Assessments to Gene Prediction and Phylogenomics. Molecular Biology and Evolution, 35(3), 543–548. 10.1093/molbev/msx319

Wick, R. R., Judd, L. M., Gorrie, C. L., & Holt, K. E. (2017). Completing bacterial genome assemblies with multiplex MinION sequencing. Microbial Genomics, 3(10). 10.1099/MGEN.0.000132

Zambrano-Mila, M. S., Agathos, S. N., & Reichardt, J. K. V. (2019). Human genetics and genomics research in Ecuador: historical survey, current state, and future directions. Human Genomics, 13(64). 10.1186/s40246-019-0249-8

